# Chemical Lithography: Selective Glyoxal Caging of mRNAs to Control Gene Expression

**DOI:** 10.64898/2026.08.25.745787

**Authors:** Alexis E. Rothchild, Devanshi C. Purohit, Jennifer M. Heemstra

## Abstract

Achieving predictable, tunable, and temporal control over mRNA function would grant direct regulation of gene expression, facilitating the development of new therapeutics and biotechnologies. Although several approaches for stimuli-responsive control of nucleic acids have been explored, most are limited to short oligonucleotides, lack a timed-release mechanism, or both. We envisioned a complementary method using glyoxal as a caging reagent. Glyoxal readily reacts with amidine groups found on the faces of nucleobases to give stable bis-hemiaminal adducts, directly disrupting hydrogen bonding. Fortuitously, this reaction is readily reversible, enabling spontaneous time-release decaging that varies with temperature. However, when applied previously to full-length mRNAs, the sequence length and excessive adduct formation resulted in no reactivation under relevant physiological conditions. To address this challenge, we developed chemical lithography in which portions of longer RNAs are “masked” through hybridization to complementary DNAs, permitting selective caging on only non-masked regions and preventing excessive adduct formation. We present an optimized glyoxalation protocol applied to EGFP as a model mRNA sequence and evaluate masking effectiveness through qualitative and quantitative studies. Using EGFP fluorescence, we monitored and assessed the ability of selective glyoxalation to control gene expression over time *in vitro*. We demonstrate the direct dependence of both the initial inhibited expression and the respective activity recovery based on the amount and location of glyoxalation. We also highlight distinct caging patterns exhibiting total inhibition upon initial treatment and complete reactivation following decaging. We anticipate that this approach will improve the mechanistic study of mRNA and gene expression and also facilitate new investigations and methods within chemical biology and biomedicine.

**Graphical Abstract:** 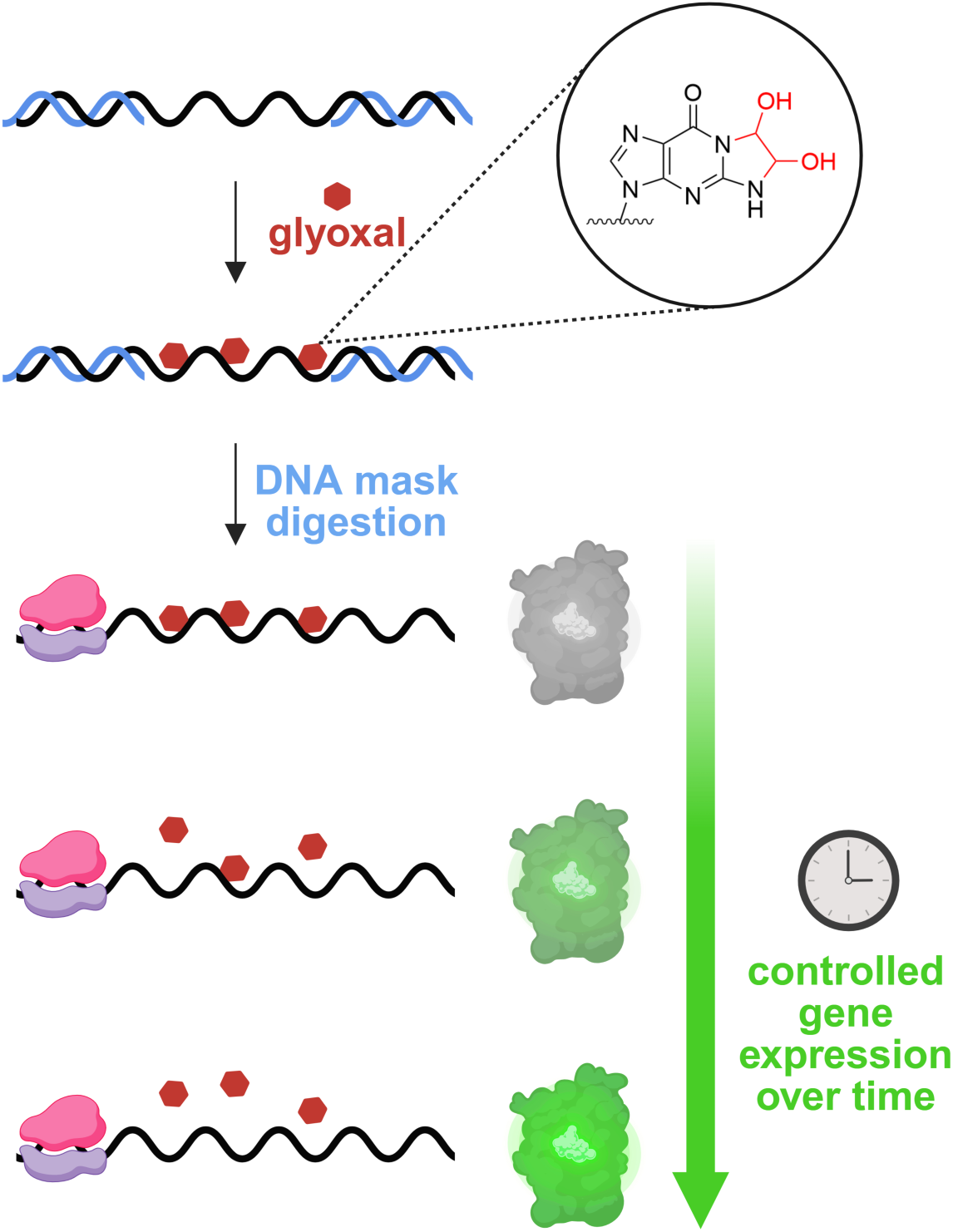

## Introduction

Nucleic acids are extremely powerful and versatile biomolecules, capable of numerous catalytic and molecular recognition activities while also playing an essential role in encoding genetic information^1^. For example, messenger RNAs (mRNAs) and the regulation of their synthesis, degradation, and availability to the translation machinery are directly responsible for the control of gene expression.^2,3^ Because of this, gaining spatiotemporal control over the functions of these molecules has become a very attractive avenue of investigation and has led to several approaches aimed at selective caging to deactivate nucleic acid function, followed by restoration at a specific time or in response to a specific stimulus.

One of the earliest and most well-known caging approaches is through the use of photoreactive monomers.^4–7^ In this method, photocleavable protecting groups, such as 9- alkoxyanthracenyl^5^ and 6-nitropiperonyloxymethyl groups,^7^ are positioned at the termini of RNAs or directly on nucleobases to “cage” the species, rendering it unable to fold, bind, or chemically react. Upon brief irradiation with light having the appropriate wavelength, the blocking groups self-cleave from the nucleic acid,^8,9^ restoring the natural base, backbone, or end structure and respective functionality. Though effective for some applications, these methods suffer from several limitations. The synthesis of these monomers is particularly difficult, and incorporation is generally only possible during chemical oligonucleotide synthesis, making longer nucleic acid targets either incompatible or prohibitively expensive to create with this approach. These approaches have also been shown to suffer from leaky caging,^8^ including incomplete inactivation and incomplete deprotection reactions, preventing full functional restoration. Some photocaging groups are also limited in compatibility within biological contexts due to the potentially damaging UV exposure and longer irradiation times necessary for cleavage.

More recently, Kool and coworkers have addressed these challenges through a “cloaking” method in which the 2’-hydroxyl groups of RNA are subject to acylation.^10,11^ This acylation effectively inactivates the oligonucleotide and the cloaking group may later be removed using either a Staudinger reduction or light-triggered decaging, depending on the specific acyl donor employed. These modifications are incorporated post-synthetically, making this approach compatible with enzymatically generated nucleic acids. Initially limited to short sequences, Kool and coworkers later expanded the approach through the development of “TRAIL” in which complementary DNAs induce reactive RNA loops, enabling selective acylation of specific regions within mRNAs.^12,13^ While offering many notable improvements over previous methods, the need for a chemical or light stimulus to induce decaging can still limit potential application space.

We envisioned a parallel approach to nucleic acid caging using glyoxal, an inexpensive and widely available reagent that can be added to and removed from nucleic acids under mild, biocompatible conditions. The reaction of glyoxal with the Watson-Crick-Franklin face of nucleobases is well-characterized and the resulting bis-hemiaminal adducts^14–16^ are known to disrupt base pairing.^17^ Guanosine adducts are the most stable and readily- formed products, and under more forcing conditions, adduct formation may also occur on adenosine and cytidine.^18^ Historically, glyoxal has been used to overcome experimental issues caused by incomplete nucleic acid denaturation^19–24^ in techniques including gel electrophoresis^22^ or northern blotting,^21^ whereby glyoxal serves as a superior denaturing agent in comparison to alternatives such as formaldehyde^24^ or formamide.^21,25^ Motivated by this unique reactivity, we recently repurposed glyoxal as a nucleic acid caging reagent,^26^ addressing many of the fundamental limitations in existing technologies. Glyoxalation is applicable to both synthetically and enzymatically generated oligonucleotides with virtually any backbone structure. More importantly, the caging reaction is reversible under mild, temperature-controlled conditions, as the adducts are stable at room temperature but undergo spontaneous hydrolysis at physiological pH and temperature.^26^ This decaging mechanism enables a unique, timed-release activation of nucleic acids under biologically relevant conditions and has already been applied to temporally regulate nucleic acid function in various contexts.^27–29^

Despite these recent advances, this caging technique has been limited in that caging occurs indiscriminately over the entire oligonucleotide sequence. This poses challenges in achieving reactivation of long oligonucleotides such as mRNA, as their length results in excessive glyoxalation, making complete decaging statistically unlikely over the typical timeframe of a biological experiment. We envisioned that a method allowing selective caging on a portion of a longer sequence would overcome this challenge by still blocking overall function but reducing the number of decaging reactions required to restore function. Excitingly, this would open the door to temporal control over mRNA expression, and we hypothesized that the location and size of the caged portion would further enable tuning of the rate of functional reactivation.

Herein, we report a strategy that allows for site-specific glyoxalation and timed-release reactivation of mRNAs. This approach, chemical lithography, utilizes complementary masking DNA oligonucleotides that hybridize to specific portions of an RNA molecule not intended to be caged, effectively blocking them from glyoxalation while leaving open stretches of RNA able to react. These DNA masks can be adjusted in length and position along the RNA, allowing for a high degree of control over where glyoxal is incorporated. Following glyoxalation, the DNA masks are removed with DNA-specific digestion treatment and subsequent purification, leaving only the selectively caged RNA. The caging can then be reversed by heating to physiological temperature or above, with the specific temperature tuning the reactivation rate. We demonstrate the ability of this method to deactivate and reactivate translation of entire mRNAs by caging specific sites on an EGFP mRNA sequence. To quantitatively monitor the efficacy of caging and kinetics of reactivation, we use a cell-free expression system^30,31^ and detect the evolution of EGFP fluorescence upon gentle heating. Together, our chemical lithography method offers a unique approach to selectively caging the Watson-Crick-Franklin face of nucleic acids and controlling gene function in a timed-release manner. We envision this will find utility in RNA-based therapeutics and other biotechnology applications.

## Results and Discussion

### Identifying conditions for selective glyoxalation of non-masked segments

Our initial report of using glyoxal to control nucleic acid function used forcing conditions to enable rapid and complete caging of the target nucleic acids. However, we recognized that these conditions would likely result in unwanted caging of masked regions in our chemical lithography approach (Figure 1a), and thus more gentle conditions to promote selective caging were needed. Specifically, these optimized reaction conditions should induce sufficient caging on non-masked RNA to achieve functional deactivation while at the same time allowing the masked regions to remain protected from adduct formation by maintaining stable RNA-DNA hybridizaiton.^32,33^ To identify these optimized conditions, we used a short, model 150 nt RNA sequence taken from the open reading frame (ORF) region of the *eGFP* gene.^34^ The sequence was left either non-masked or hybridized to three 50 nt complementary DNA masks for initial testing (Figure 1b), enabling us to search for reaction conditions that would lead to the desired selectivity.

**Figure 1.**
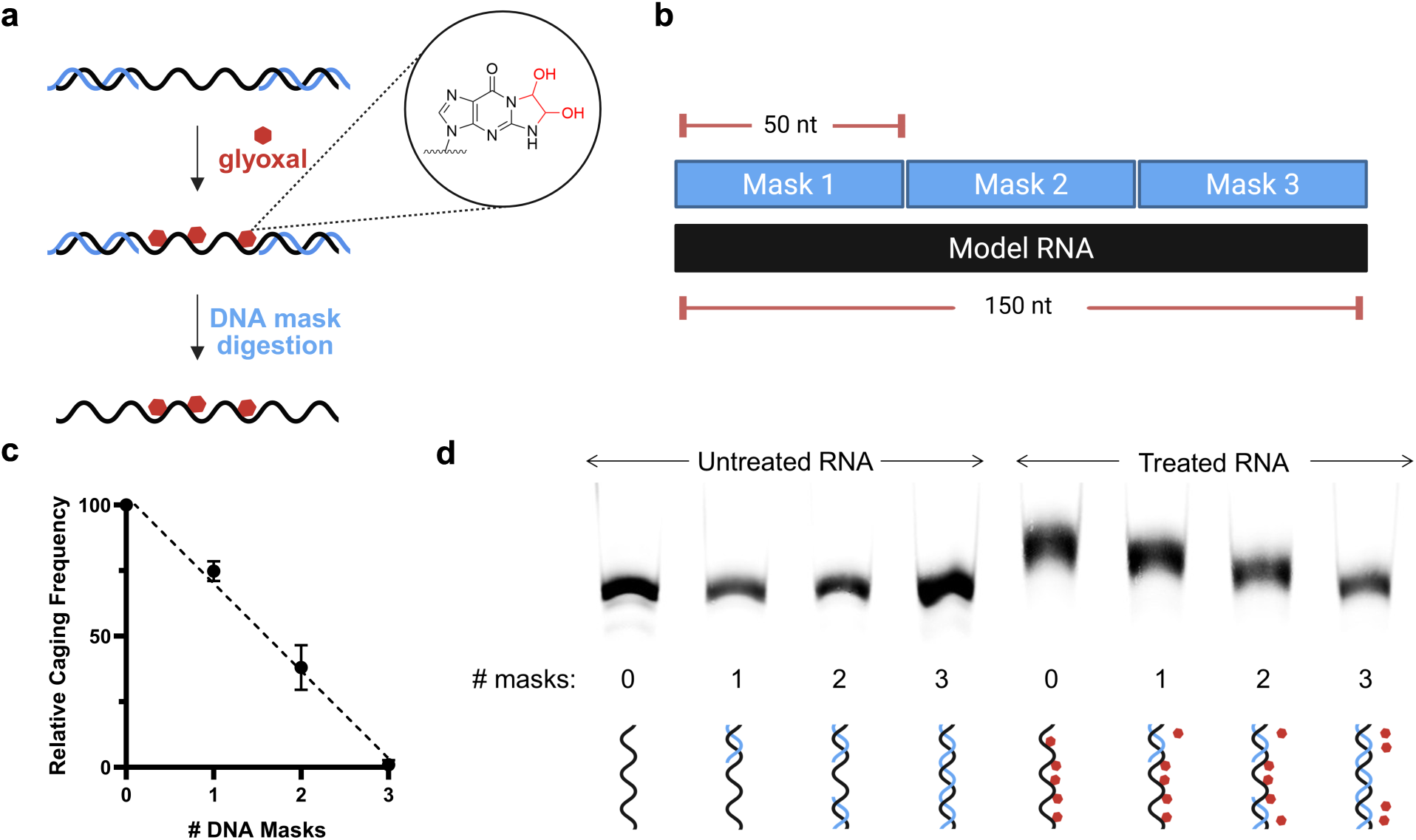
Glyoxal caging optimization. (a) Overview of chemical lithography. (b) Model RNA design used during optimization. (c) Calculated caging frequency of each masking design. Experiment was performed in triplicate and error bars represent the standard deviation. (d) Analysis of caging of model RNAs having varied masked regions using 10% denaturing PAGE. All RNAs were reacted with 100 mM glyoxal at 30°C for 1 h in PBS with 1 M added NaCl. All RNAs are Cy5-labeled.

Using the previously established reaction conditions for full caging of an RNA (1.3 M glyoxal in 50% DMSO, 50°C 1 h),^26,35,36^ we systematically altered each parameter to achieve the desired specificity. We first changed the reaction buffer from DMSO to PBS (Figure S1), avoiding the potential hydrogen bond disruption induced by the organic solvent.^37–39^ We then individually varied glyoxal concentration, reaction temperature and time, and the addition of salt to stabilize the masked duplexes.^40–42^ Reactions were monitored via denaturing polyacrylamide gel electrophoresis (PAGE), with relative caging extent being observed as an apparent shift in molecular weight (Figure S2a-d).^26^ Each optimized parameter was chosen by identifying the conditions that generated a visible band shift for non-masked RNAs while maintaining no detectable band shift for fully-masked RNAs. After analyzing each parameter, we arrived at final conditions of 100 mM glyoxal at 30°C for 1 hour in 5x PBS with 1 M added NaCl. To fully evaluate the efficacy of these reaction conditions, we assessed the relative adduct formation on the 150 nt RNA under varying masking conditions through denaturing PAGE (Figure 1d). Excitingly, fully-masked RNAs consistently displayed no caging, indicated by the lack of band shift. When single masks were removed from the reaction, the RNA showed increasing amounts of caging as would be expected (Figure 1c). These results offer convincing evidence that our optimized reaction conditions allow for efficient caging of non-masked RNA yet remain mild enough for complementary DNA masks to block caging.

### Validating selectivity of chemical lithography

Analysis of glyoxalation by PAGE provided encouraging data to suggest we had achieved selective reaction of the non-masked segments, and we sought to further validate this through assessment of site-specific impact on RNA hybridization. Using a 3’ fluorescently end-labeled 150 nt RNA, we designed a complementary 50 nt DNA having a 5’ quencher (Q-DNA) that would be positioned in close proximity to the fluorophore if hybridization were to occur (Figure 2a). We confirmed that a non-quencher-modified DNA mask did not lead to detectable quenching (Figure S3b), then tested increasing concentrations of Q- DNA to quantify the maximum fluorescence quenching possible upon hybridization to untreated, fluorescently-labeled RNA (Figure S5a, b)

**Figure 2.**
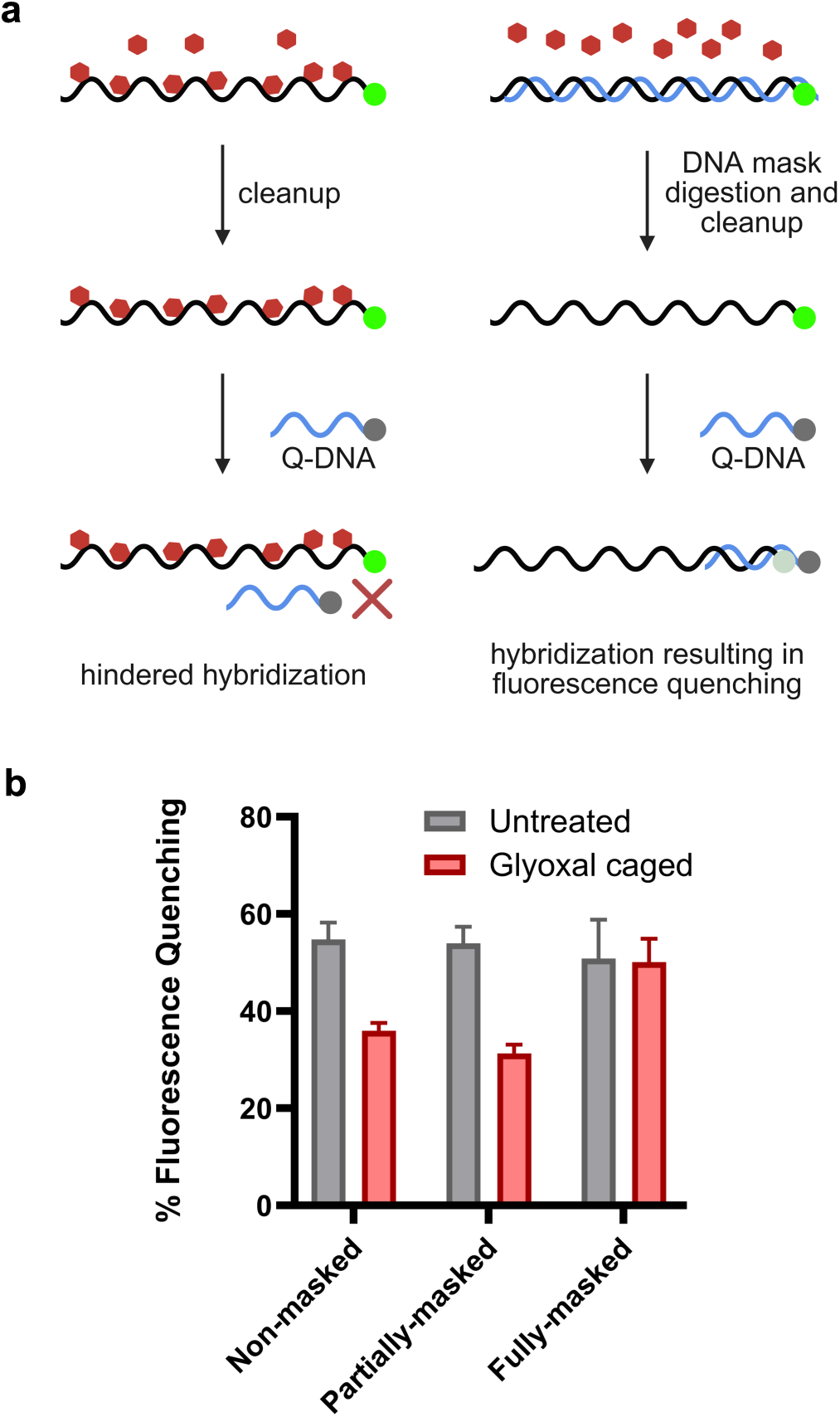
Q-DNA assay. (a) Q-DNA assay workflow. (b) Normalized quenching for differentially masked model RNAs with and without exposure to glyoxal. The % fluorescence quenching represents the ratio of the fluorescence intensity for RNA with Q-DNA versus RNA alone. Each experiment was run in triplicate and error bars represent the standard deviation.

Interestingly, during these preliminary studies we observed an unanticipated quenching upon exposure of some fluorophores to glyoxal. In particular, Cy5-labeled RNA exhibited significantly reduced signal following glyoxal treatment, independent of the presence of the Q-DNA. This effect is likely due to a reaction between glyoxal and the amine groups present on this fluorophore, causing a structural change and resulting in reduced fluorescence. To overcome this issue, we systematically analyzed the effect of glyoxal treatment on a panel of fluorophores (Figure S4) and found that FTSC (fluorescein-5-thiosemicarbazide) retained robust fluorescence following exposure to 100 mM glyoxal, indicating compatibility with our chemical lithography approach.

With an appropriate fluorophore identified, samples having varying masking and caging patterns were evaluated. We compared samples that were non-masked, fully-masked, or partially-masked over the first 100 nt, such that the fully- and partially-masked samples differed in the presence or absence of masking over the region where the Q-DNA was designed to bind. Each sample was assessed both before and after glyoxal caging and subsequent mask digestion, and the resulting levels of fluorescence quenching are shown in Figure 2b. The % fluorescence quenching was calculated as the ratio of signal between samples incubated with and without the Q-DNA strand. This normalized quenching was individually measured for each differently-masked RNA and can be equated to relative binding efficacy of the Q-DNA across these samples. As anticipated, the non-masked and partially-masked samples showed a significant decrease in % fluorescence quenching after treatment with glyoxal, as glyoxalated adducts were able to form in the Q-DNA hybridization region and prevent binding of the quencher-labeled DNA. Excitingly, in the case of the fully-masked sample, we observed no change in % fluorescence quenching between treated and untreated samples, indicating that the DNA mask was able to prevent glyoxalation. Under more forcing conditions (1.3 M glyoxal, 50°C, 4 h), the level of fluorescence quenching was further reduced (Figure S6), suggesting that the partial quenching observed for the non- and partially-masked samples may arise from the ability of the Q-DNA to still bind weakly to a more sparsely caged sequence. However, this is not of concern for our primary application of modulating protein expression, as we expect that even a single glyoxal adduct is sufficient to stall translation.

### Optimization of DNA mask removal

Before exploring the effects of selective glyoxalation on mRNA function, we recognized that attention was also needed toward optimizing the DNA digestion workflow. Incomplete DNA mask digestion and removal could result in unintended mRNA silencing induced by antisense effects,^43,44^ and thus optimizing this step was crucial for use of our selective caging strategy to modulate gene expression. We utilized full-length EGFP mRNA as our model mRNA to test for DNA mask removal and monitored the ability to achieve *in vitro* EGFP expression, quantified as RFU/hr, in comparison to non-masked samples. We tested various nucleases (Figure S7a), reaction conditions, (Figure S7b,c), purification methods (Figure S7a), post-digestion denaturants and clean-up columns (Figure S7d), as well as the order in which these steps occur (Figure S7e). We found an optimized workflow in the form of a sequential DNA digestion with RNA-specific purification methods following each enzymatic reaction (Figure S7e,), which resulted in the highest translation efficiency of all methods tested. Without the optimized digestion and purification workflow, mRNA silencing proved to be an extremely potent effect, causing a complete loss in signal when contaminating DNA remained (Figure S8).

### Gene expression of selectively caged mRNAs

With an efficient mask removal workflow elucidated, we were ready to test the ability of our chemical lithography approach to temporally modulate gene expression. To achieve precise and controlled measurements, we utilized the EGFP mRNA in a cell-free gene expression system and performed experiments in parallel for samples having selective glyoxalation at multiple specific regions on the mRNA (Figure 3a). As with our model system, we hybridized portions of the mRNA not intended to be caged with DNA masks, leaving the open regions subject to glyoxalation. The DNA masks, 60-85 nt in length, were designed to completely tile the mRNA, and omission or modification of one or more masks enabled the specific regions of interest to remain non-masked and thus receptive to glyoxalation. We tested ten different glyoxalation patterns, varying in both length and location along the mRNA. Longer portions included the entirety of the 5’ or 3’UTR regions as well as the complete ORF split into upstream and downstream regions. Within these longer sites, we also tested two varying designs of the 3’UTR region: one open region beginning immediately after the stop codon (3UTR V1) and the second open region encompassing only the poly(A) tail (3UTR V2). Shorter sites investigated include the Kozak Sequence (KS), start codon sequence (C1), random codon sequences in both the upstream (C10) and downstream ORF (C212), corresponding to codon 10 and codon 212 respectively, and the sequence of the last three codons in the ORF (C236-238), including the stop codon. A detailed gene map of these sites can be found in Figure 3c. All indicated regions were caged using the optimized glyoxalation conditions followed by DNA mask removal using our optimized digestion workflow. For each differentially masked sequence, we compared glyoxal treated with untreated control, enabling us to calculate the normalized gene expression and control for any lost signal due to digestion and purification steps (Figure 3d,e). For these gene translation studies, we also used fully-masked and completely non-masked EGFP mRNAs as positive and negative controls, respectively, to compare the effect of glyoxalation across all masking patterns (Figure 3b). The fully-masked samples consistently displayed minimal to no inhibition when treated with glyoxal, further validating the efficacy of our masking approach and providing us with a reliable control to compare to the selectively caged RNAs.

**Figure 3.**
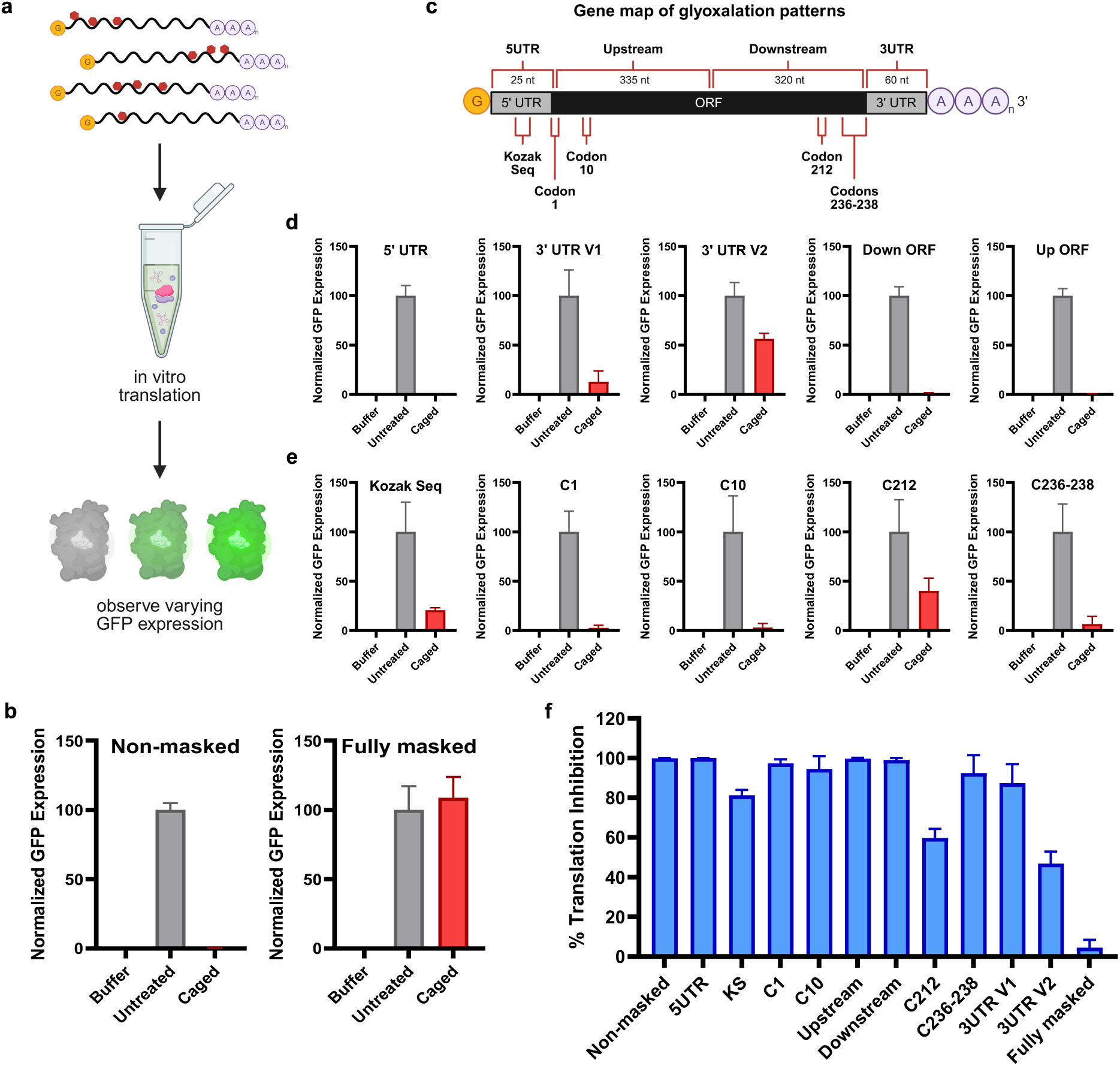
*In vitro* gene expression of selectively caged mRNAs. (a) *In vitro* translation workflow. (b) GFP expression of completely non-masked and fully-masked EGFP mRNA, serving as both the negative and positive controls. (c) Gene map of regions of interest. “Codon #” denotes which codon corresponds to each respective RNA sequence targeted. (d) Normalized GFP expression of longer caged sites. (e) Normalized GFP expression of shorter caged sites. (f) Relative translation inhibition of all sites when caged with glyoxal. Each experiment was run in triplicate and error bars represent the standard deviation.

Excitingly, the differentially masked mRNAs displayed varying GFP expression depending upon the size and location of the caging window. Longer caged regions, both upstream and downstream in the ORF, completely inhibited translation when compared to untreated controls. This is expected due to the large number of adducts readily formed on these longer portions, presumably causing ribosome stalling. Caging of the 5’ UTR also completely inhibited translation, likely through a combination of stalling the ribosome while scanning for the Kozak Sequence (GCCACCAUGG) and direct caging of the Kozak sequence, preventing translation initiation. The 3’UTR caged mRNAs showed different changes in expression when glyoxalated, with V1 retaining 15% of original activity and V2 retaining 55% of original activity. This suggests that caging directly following the stop codon may hinder the ribosome from efficiently completing translation. Conversely, caging in the poly(A) tail only minimally inhibits gene expression and this slight inhibition seen on V2 could be due to downstream destabilization effects caused by the glyoxal adducts formed along the poly(A) tail.^45^

We observed the greatest modulation in expression levels upon glyoxalation for the smaller caged regions. For sites that are directly responsible for ribosome recognition and translation initiation, KS (GCCACC exposed) and C1 (Met1^AUG^ exposed), robust inhibition of gene expression was observed, with C1 specifically showing complete blockage of any detectable translation. C10 and C236-238 displayed similar expression inhibition to these regions as well. C10 is comprised entirely of guanine residues (Gly10^GGG^ exposed) and modification of any of these would likely be sufficient to induce ribosome stalling and premature truncation of the nascent peptide. A caged stop codon and three terminal amino acid codons (LeuTyrLys236-238^CUG^ ^UAC^ ^AAG^ ^UGA^ exposed) on the other end of the ORF also inhibits translation and EGFP maturation to the same degree. For C212, significantly less inhibition was seen, which is expected given that this codon contains no guanines (Asn212^AAC^ exposed) and thus likely does not achieve full caging. Together, these results demonstrate the efficacy of chemical lithography, in that gene expression can not only be turned off, but that the level of attenuation can be modulated by varying the region of the mRNA that is subjected to caging.

### Translation reactivation as a function of caging window

We were very encouraged by our ability to achieve varying levels of translation inhibition using different mRNA caging windows and hypothesized that time-dependent reactivation of translation would be similarly tunable. In choosing parameters for these experiments, we sought to mimic physiological conditions as closely as possible, but also needed to balance this with the fact that glyoxal decaging typically has a half-life of days at 37°C and thus requires monitoring for at least 7 days to observe maximum decaging, but mRNA degradation becomes a challenge when this long of timeframe is used for *in vitro* translation experiments. Fortunately, we found that the wheat germ extract buffer used for *in vitro* translation could be adjusted to physiological pH without impeding translation (Figure S9).^26^ For temperature, we decided to perform experiments at 50°C, as this remains rather close to physiological conditions but allows for decaging to occur before RNA degradation or sample evaporation become problematic. Importantly, the purpose of these experiments was to establish our ability to tune the reactivation rate for mRNAs depending on their caging pattern, and we would expect these trends to be consistent across a range of decaging temperatures. Further, we have previously established the ability of shorter RNAs to undergo time-dependent reactivation in living cells at 37°C over the course of 7 days. Thus, the insights gained in these experiments can be directly applied to control mRNA reactivation in biological systems where these limitations of cell-free systems are no longer of concern.

As shown in Figure 4a, our workflow consisted of translating the selectively caged mRNAs at pre-determined timepoints during the decaging incubation. We monitored activity at six different decaging timepoints from 4-72 h, using the EGFP expression recorded upon initial caging as the 0 h data point, and quantified gene activity using the *in vitro* translation assay. For every glyoxal treated sample, we used a control sample that had undergone the same DNA masking and unmasking protocol, but had not been treated with glyoxal, and monitored EGFP expression at the same timepoints. This enabled us to calculate activity recovery for each caging pattern across each of the timepoints to determine the effect of the caging window and size on reactivation rate.

**Figure 4.**
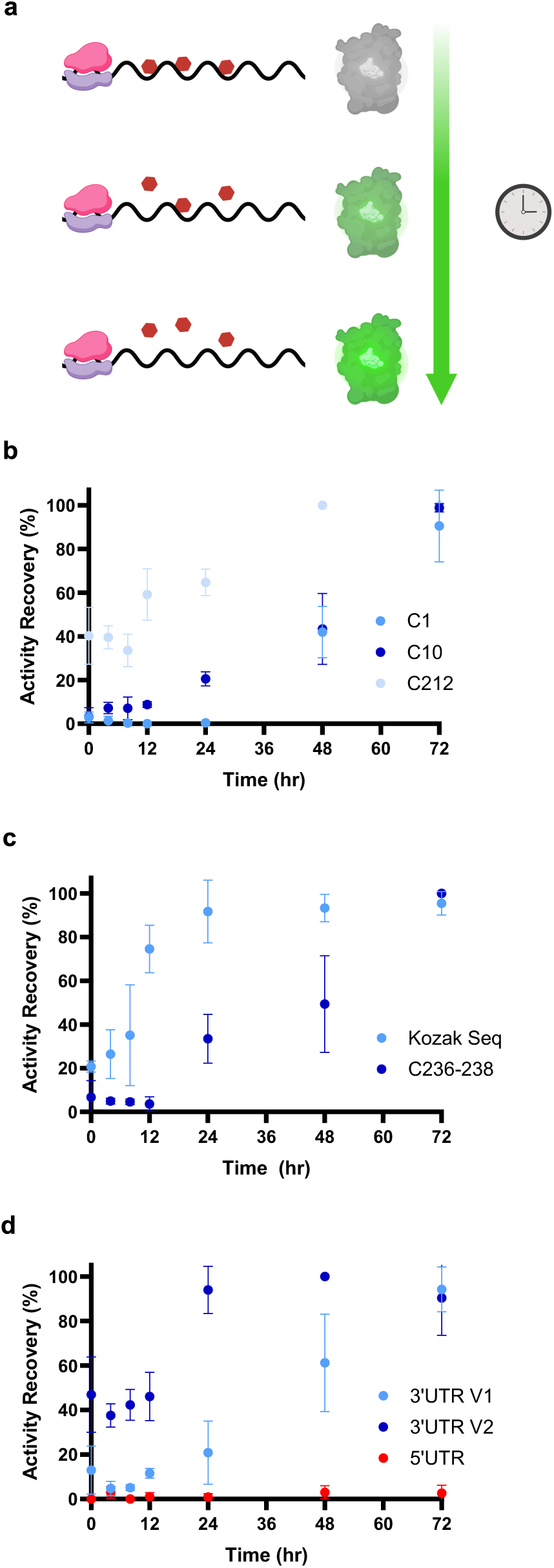
Decaging and recovery of *in vitro* GFP expression for selectively caged mRNAs. (a) Representation of decaging inducing increased gene expression over time. (b) Activity recovery of single codon caged sites. (c) Activity recovery of shorter caged sites. (d) Activity recovery of UTR-caged sites. All experiments were performed in triplicate at 50°C and error bars represent the standard deviation.

Excitingly, we achieved complete reactivation for seven of our ten mRNA caging patterns. These designs include all of the shorter selected regions and both the 3’UTR-caged variants. For the single codon caged samples, total reactivation was observed after 48- 72 h of incubation (Figure 4b). This includes even C1, in which the start codon is left non-masked and where we observed complete inhibition of gene expression when caged. This capability places our chemical lithography method in a unique application space with being the only method that allows full-length mRNA to have translation completely blocked by caging and then fully restored in a spontaneous, time-release fashion. Additional caging patterns targeting shorter portions of the mRNA, such as the Kozak Sequence and the final three codons of the ORF, also displayed complete reactivation after 72 h of incubation. (Figure 4c). The sample caged along the Kozak Sequence (GCCACC) achieved full restoration after only 24 h of incubation, representing the fastest sample to reactivate translation. We expect that only the single guanine in this sequence undergoes caging, though that still leaves an interesting question as to why reactivation is faster than other caging patterns, such as C1, that also contain a single guanine. One potential hypothesis is that even with the guanine caged, the Kozak Sequence is still able to recruit the small ribosomal subunit, which then participates in accelerating the decaging reaction to allow for assembly of the full ribosome.^46–48^ In future work, we would be interested to test mRNAs having a GCCGCC Kozak Sequence to determine whether reactivation rate can be further tuned by the presence of a second guanine nucleotide that can be caged. The last of our mRNAs exhibiting complete activity restoration are the two caging patterns focused on the 3’UTR (Figure 4d). V2, having only the poly(A) tail non-masked, reactivated much faster at ∼24 h of incubation whereas V1, having a longer, non-masked region, required 72 h, which is consistent with their expected caging patterns.

For the remaining three mRNA caging patterns, no restoration of translation activity was observed, which was not unexpected. The caged regions encompassing the entirety of the upstream and downstream ORF proved too long to achieve timely quantitative decaging, showing no detectable EGFP expression even after 72 h (Figure S11). Similarly, the 5’UTR caged region exhibited no detectable recovery following the same incubation time (Figure 4d). Though having a significantly shorter caging window than the ORF designs, we hypothesize that the high capacity for adduct formation on this 25 nt window combined with the essential role it plays in ribosome recognition, together prevent detectable reactivation. These results further underscore the importance of our chemical lithography approach for limiting caging to specific sections of an mRNA such that activity can be restored over the desired time period.

## Conclusion

Glyoxal caging provides a powerful approach to modulate the function of nucleic acids and, compared to typical methods that involve removal of caging groups using a chemical or light stimulus, glyoxal undergoes spontaneous timed-release decaging under physiological conditions. While glyoxal has been shown to be highly effective in temporarily caging aptamers, antisense therapeutics, and other short oligonucleotides, we have found that this approach does not directly extend to longer nucleic acids such as mRNAs. Here we overcome this challenge using chemical lithography, in which DNA masks are employed to block specific sections of the RNA, allowing caging to selectively occur in only the non-masked regions and thus creating highly specific caging patterns even on large RNAs.

We optimized reaction conditions to provide robust caging in non-masked regions while preventing caging in masked regions and validated this selectivity using both qualitative and quantitative assays. We also optimized conditions for removal of the DNA masks to enable translation of the resulting mRNA. To demonstrate the utility of our chemical lithography method, we explored application in modulating *in vitro* translation of selectively masked and caged EGFP mRNA. We found that the size and location of the caging window directly impact the resulting inhibition of EGFP expression upon caging as well as the rate of spontaneous reactivation. Larger exposed regions completely inhibited gene expression upon caging but were resistant to decaging efforts, similar to our experiences caging the full mRNA sequence. Excitingly, motifs having smaller exposed regions targeted for their role in ribosomal recognition or guanine content, were found to effectively inhibit gene expression upon caging and were successfully reactivated upon incubation over hours to days in mild, biologically relevant conditions. This demonstrates that chemical lithography enables the caging and time-dependent reactivation of even long RNAs, and that the specific rate of reactivation can be controlled by modulating the size and location of the caging window. We anticipate that this approach will have utility in basic science applications such as controlling synthetic biology gene circuits, as well as biomedical applications such as timed-release formulations of mRNA therapeutics.

## Associated Content

### Supporting information

Experimental materials and methods including supplementary tables and supporting figures (PDF)

## Author Contributions

Conceptualization, J.M.H.; methodology, A.E.R., D.C.P., and J.M.H.; investigation, A.E.R.; data analysis, A.E.R; writing, A.E.R. and J.M.H.; and supervision and funding, J.M.H.

## Notes

The authors declare no competing financial interest.

## Acknowledgements

This work was supported by the National Science Foundation (CHE 2306047 to J.M.H.) and the National Institutes of Health (R35GM144075 to J.M.H.). This content is solely the responsibility of the authors and does not necessarily reflect the official views of the National Science Foundation. Schematics were made using BioRender.com. We would like to thank Drs. Steve Knutson, Aimee Sandford, Alexandria Quillin, and Tyson Todd for helpful conversations, mentorship, and advice and Tyson Todd for experimental guidance and assistance in drafting the manuscript.

